# The impact of a reduced calorie, macronutrient diet change on Caucasian males in relation to genotypes associated with obesity, increased BMI and dietary response

**DOI:** 10.1101/425702

**Authors:** L Ellis, C. Collins

## Abstract

Obesity is a growing crisis within the 1st world and the causes of obesity are well studied: a lack of physical activity combined with an increased calorie intake. This energy imbalance leads to fat accumulation leading to eventual obesity. Whilst the cause may appear clear, the options of lifestyle change and/or diet planning that is efficient with effective sustainability has been harder to come by. Genetic variants linked to obesity may hold clues on how to best customise a diet plan to an individual leading to an easier to change lifestyle with better sustainability. This study aimed to find correlations with a diet change that decreased kcal intake by 200kcal per day with a macronutrient alteration (decreased carbohydrates and increased unsaturated fat) and genetic components to discover responders and non-responders. Within the study group of Caucasian males the genes TCF7L2, TFAP2B, PPARG2 and FTO all impacted the response of the dietary intervention. In conclusion it would be prudent to consider DNA analysis as part of a battery of diet and lifestyle tests to personalise a sustainable and effective lifestyle/dietary intervention to help control weight.

## Introduction

Common knowledge of obesity is a result of when energy intake chronically exceeds energy expenditure, creating a positive net energy balance. And even though the prevention and treatment of obesity is becoming a better studied topic obesity is still increasing worldwide. The fact is, the pathophysiology of the condition is complex and the causes of energy intake and energy expenditure are multifactorial. The main methods of energy expenditure are:

1. BMR
2. Thermic effect of food (TEF)
3. Physical activity
4. Non-exercise activity thermogenesis (NEAT)
5. Newly found genetic predispositions which can increase or decrease natural energy expenditure.

Data has shown that there is a genetic contribution to obesity within populations. Individuals are more likely to develop the phenotypic expression of obesity based upon a permissive role of the environment with the main link being a strong genetic predisposition. Over 40 sites in the human genome have been discovered that can increase genetic susceptibility to obesity including polygenic cases of obesity; multiple gene mutations, single nucleotide polymorphisms and Mendelian obesity ^(1)^

Due to this, nutritional genomics suggests that individuals in the same obesogenic behaviour (environmental interactions), those with genetic mutations that promote obesity will likely gain weight, whereas those without the mutations won’t gain weight. Obesity should not be treated as a phenotypic disease; instead effective treatment should incorporate the combination of environmental, physiological, psychological, biochemical and genetic factors ^(2)^.

A report outlined that 127 genes with one positive association to obesity had been found and among these, 22 were supported by five positive studies ^(3)^. These genes were: ACE, ADIPOQ, ADRB2, ADRB3, DRD2, GNB3, HTR2C, IL6, INS, LDLR, LEP, LEPR, LIPE, MC4R, NR3C1, PPARG, RETN, TNFA, UCP1, UCP2, UCP3 and VDR.

Studies suggest there are five overall genotypes, to which numerous genes can be classified into. These genotypes are:

1. The thrifty genotype – in this genotype, the individual has a low metabolic rate and therefore lower levels of thermogenesis.
2. The hyperphagic genotype – these individuals have poor control over appetite and low satiety levels which means they have an increased risk of overeating
3. The sedentary genotype – this genotype is linked to those with low levels of physical activity (above normal NEAT)
4. The lipid oxidation genotype – these individuals have low levels of lipid oxidation, therefore they do not use fat stores as an energy source in an efficient manner.
5. The adipogenesis genotype – these genotypes are associated with a higher lipid storage capacity which means that overfeeding is likely to expand the adipocyte storage.

There is a synergistic relationship between environmental factors and genes which means that when genetic mutations are present, the risk of obesity is increased and the severity of which is determined through interactions with the environment.

## Methods

**The aim of this study** was to: To investigate the association between genotype and BMI in Caucasian males (n=92) aged between 29 and 38.

Using genetic screening as a tool to improve obesity treatment is becoming more prevalent. The aim of this study was to assess the association between genotypes, BMI and weight. Further to this, macronutrient alteration was used by changing KCAL from carbohydrate to KCAL from fats along with a simple reduction of 200kcal per day over a period of 24 weeks. BMI and weight were assessed in active healthy males (n=92) to determine association of genotypes using a quasi-experimental study design. 78 genes were initially assessed during the 24 week programme that altered only dietary factors of the individuals’ lifestyle, no exercise or physical activity protocol were introduced:

- Alteration to macronutrients, 350kcal (87.5 - 90g) was cut from predominately sugar based foods and replaced with 150kcal (16.6 - 20g) of predominately unsaturated fat based foods, leaving a calorie deficit of only 200kcal per day.

### Outliers

Values for total weight loss and BMI loss were checked for unrealistic values and outliers and such values and outliers were excluded from the assessment. Values of weight loss >=15kg were excluded as they were deemed too excessive following the 24 week intervention.

### Stats test

Of the 78 genes which were assessed, only 8 drew attention after removing outliers and removing any missing genotypes. A Hardy-Weinberg test was conducted on all 8 genotypes to show equilibrium. T-tests were conducted for baseline BMI before and after the 24 week intervention. Significant genes from the T-tests underwent a linear regression model to analyse the effects of genotypes on BMI and weight.

## Results

5 participants were excluded from the study due to weight loss equal to or above 15kg which was deemed excessive for 24 weeks.

Participants had a mean baseline BMI of 27.9 kg/m^2^ and mean weight was 90kg at the start of the study, Following the macronutrient alteration to the diet with a decrease in carbohydrate intake and increase in fat intake on average participants lost −8kg and their BMI decreased by 3.5 kg/m^2^. (Table2).

**Table 1:**
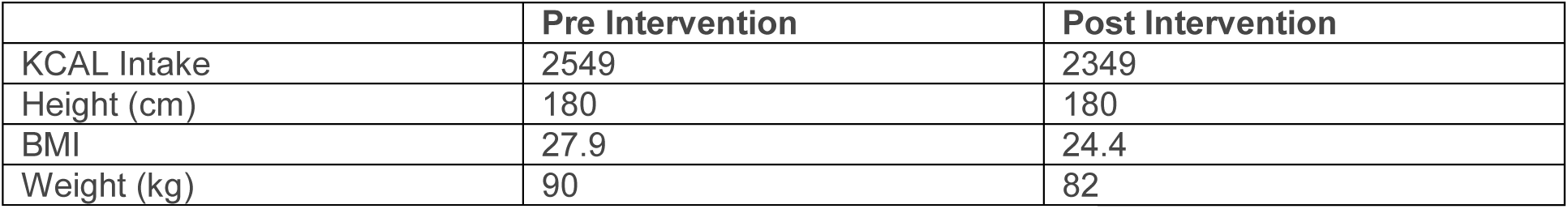
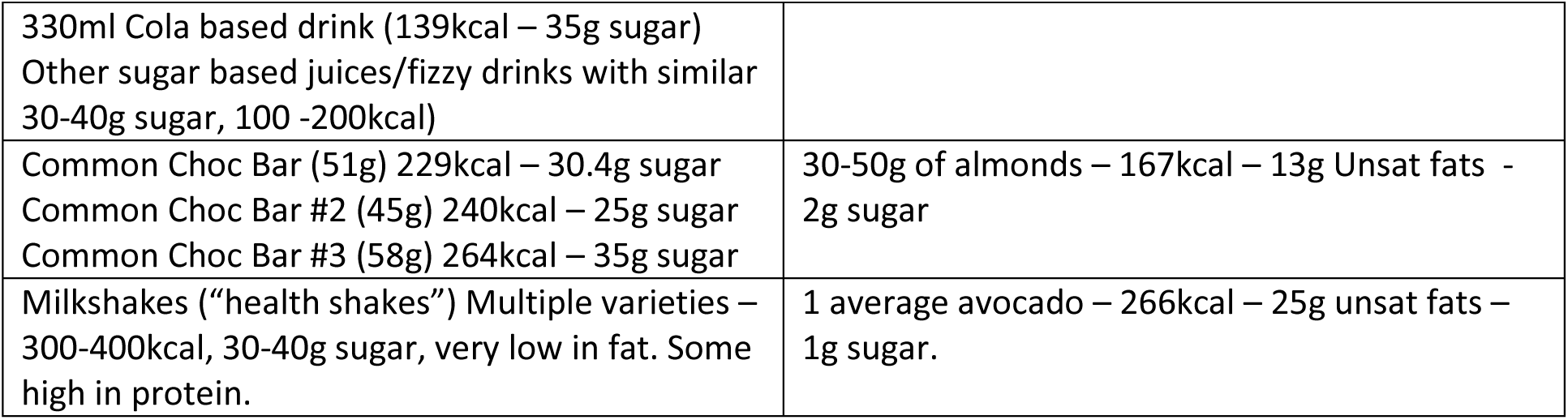
Example of food swaps

**Table 2:**
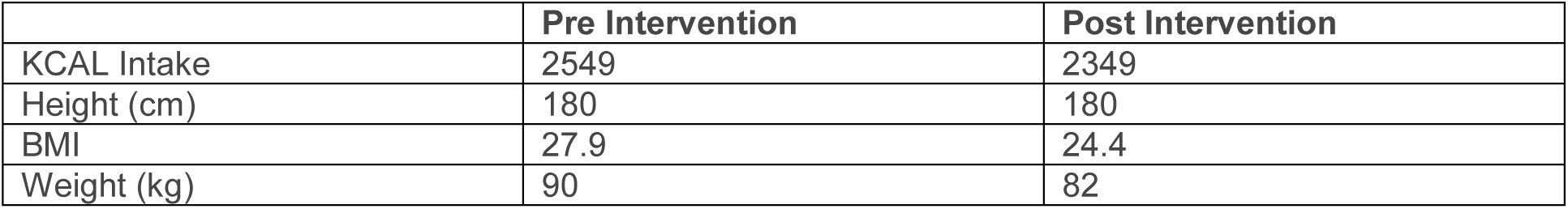
Descriptive statistics of average pre and post intervention.

8 genes and their respective polymorphism were put into the Hardy Weinberg equilibrium (Table 3) and further analysis was conducted on these genes. T-tests on these 8 genes isolated the results further, and found only 4 genes to be statistically significant pre and post intervention on BMI and weight loss outcomes. A regression analysis then looked into the effects of the genotypes on the 4 genes which were shown to have significant differences. These were:

**Table 3.** Showing the four genes which were in Hardy-Weinberg equilibrium and had significant effects pre and post intervention. (^∗^NC/HEZ/HOZ = Non carriers / heterozygotes / homozygotes)

| Gene | SNP | Risk Allele | Genotype |  |  | Hardy Weinberg |  |
| --- | --- | --- | --- | --- | --- | --- | --- |
|  |  |  | NC | HEZ | HOZ | R<sup>2</sup> | P |
| TCF7L2 | rs7903146 | T | 24 | 5 | 9 | 1.35 | 0.28 |
| TFAP2B | rs987237 | G | 21 | 16 | 1 | 0.18 | 0.66 |
| PPARG2 | rs1801282 | G | 31 | 6 | 1 | 1.27 | 0.25 |
| FTO | rs9939609 | A | 12 | 22 | 4 | 0.85 | 0.35 |

### TCF7L2

Average weight loss in C allele (n=24) was −8 KG and BMI loss was −2.6 In the CT (n=5) was −6kg and −1.9. For homozygote T (n=9), no significant difference were found pre and post intervention for weight along (T(34) = 1.09, P < 0.05). No other significant effects were found. Significant differences between pre and post for C allele (T(64) = 6.7, P < 0.05) and between CT allele (t (5) = 7.79, p < 0.05) but no significant differences between groups (T(36) = 2.71, P > 0.005).

**Graph 1:**
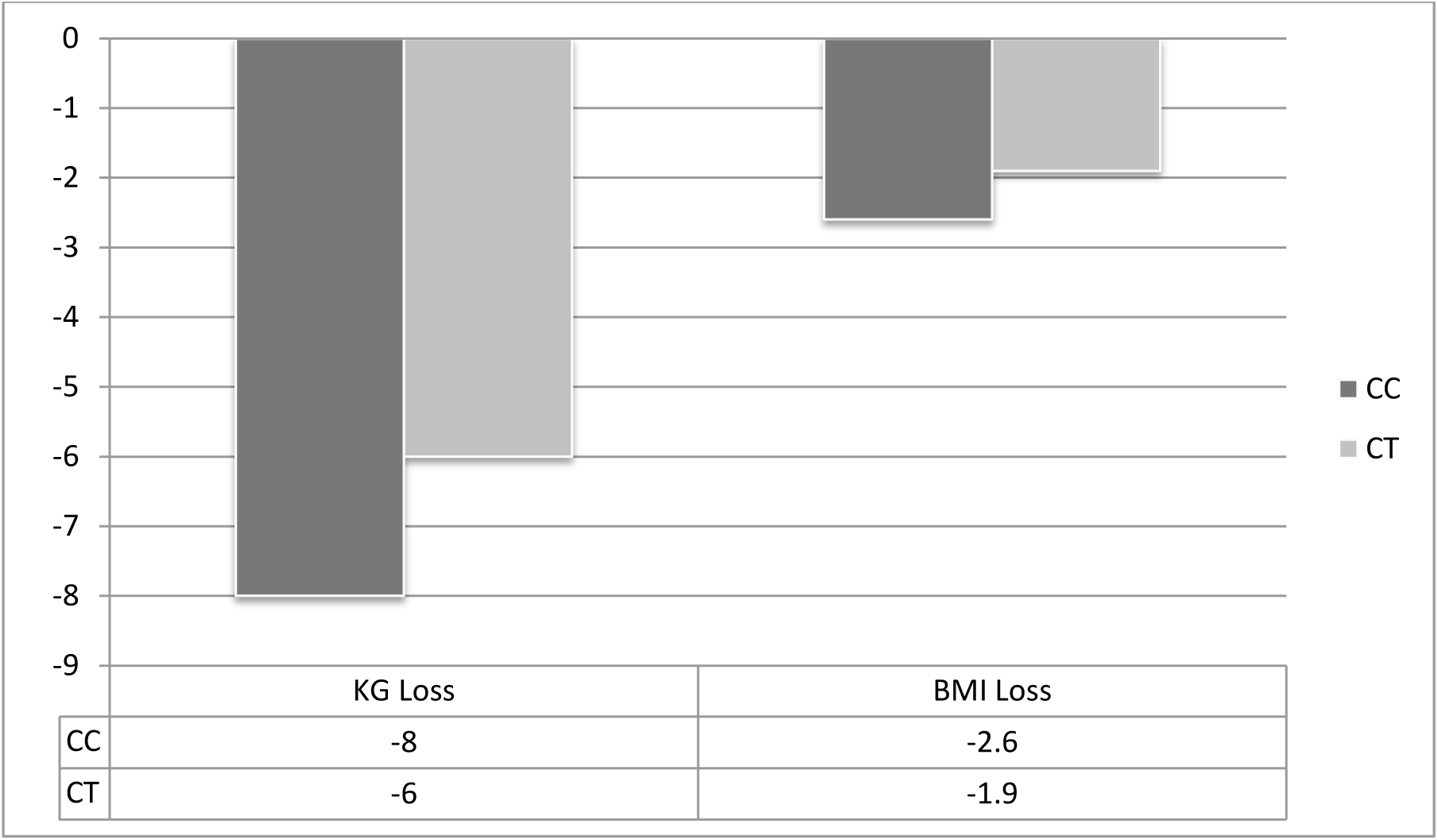
TCF7L2 gene variants, weight and BMI change pre and post intervention.

### TFAP2B

Average weight loss for A allele (n=21) was −7KG and BMI loss was −2.27 with significant difference (T (40) = 4.86, P< 0.05). Only one participant was homozygous for G allele so data was excluded. For GA allele (n=16) average weight loss was −9kg and BMI loss was −2.87 with significant difference (T (30) = 4.74, P< 0.05). A significant difference was also found between groups for A allele and GA allele, suggesting the A allele was important in controlling weight loss (T (35) = 4.66, P < 0.05).

**Graph 2:**
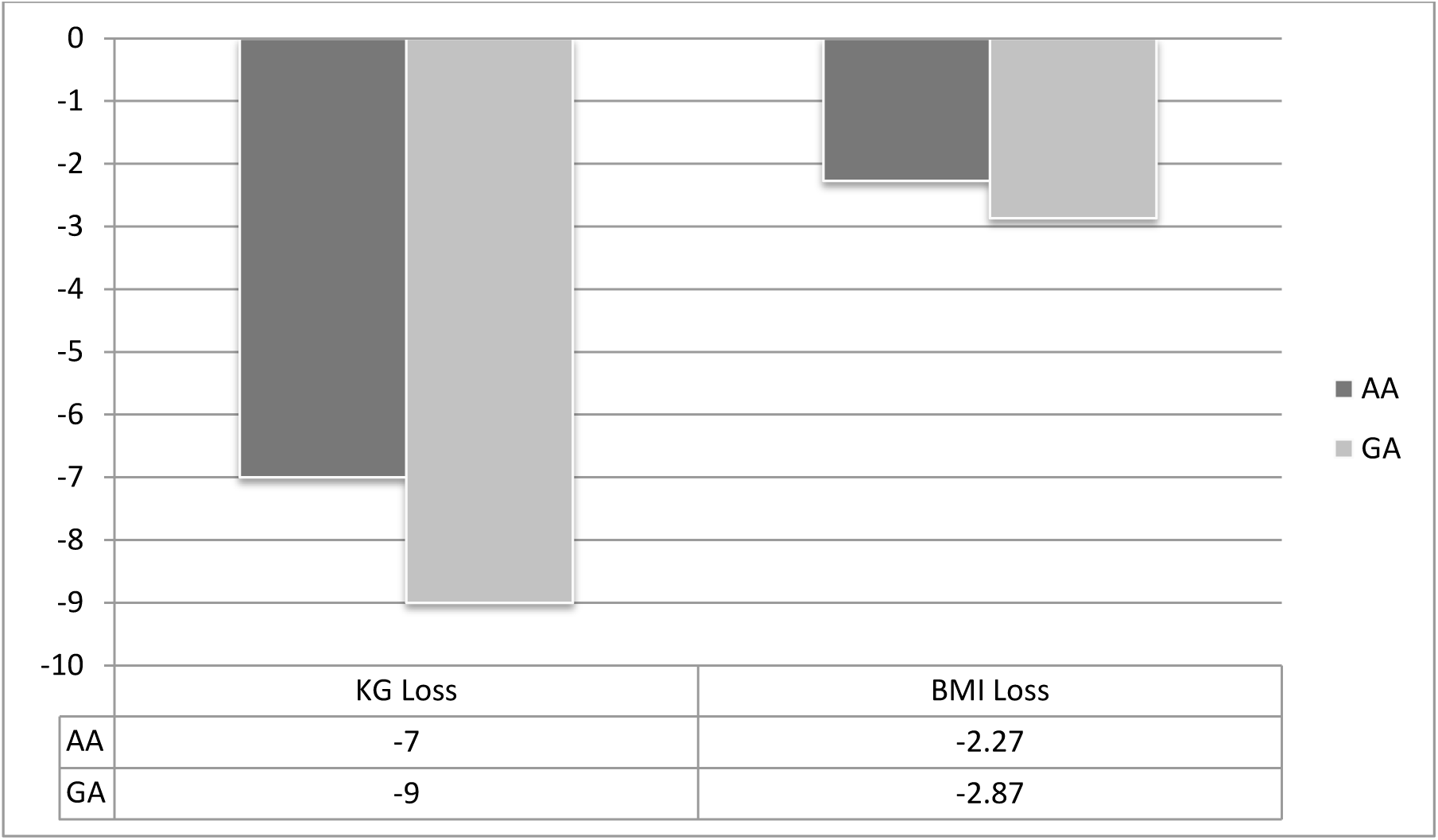
TFAP2B gene variants, weight and BMI change per and post intervention.

### PPARG2

average weight loss for C (n=31) allele was −8kg and for BMI was −2.41 with significant difference before and after intervention (T (60) = 6.12, p < 0.05). Only one participant was homozygous for G allele so data was excluded. For GC individuals (n = 6), average weight loss was −10kg and BMI loss was −3.23 which was the highest BMI loss out of all 8 genotypes identified to be significant. Significant differences was found pre ad post intervention for BMI for GC alleles (T (10) = 3.23, p < 0.05) and similarly pre and post between C homozygous and GC heterozygotes (T (35) = 4.77, P < 0.05).

**Graph 3:**
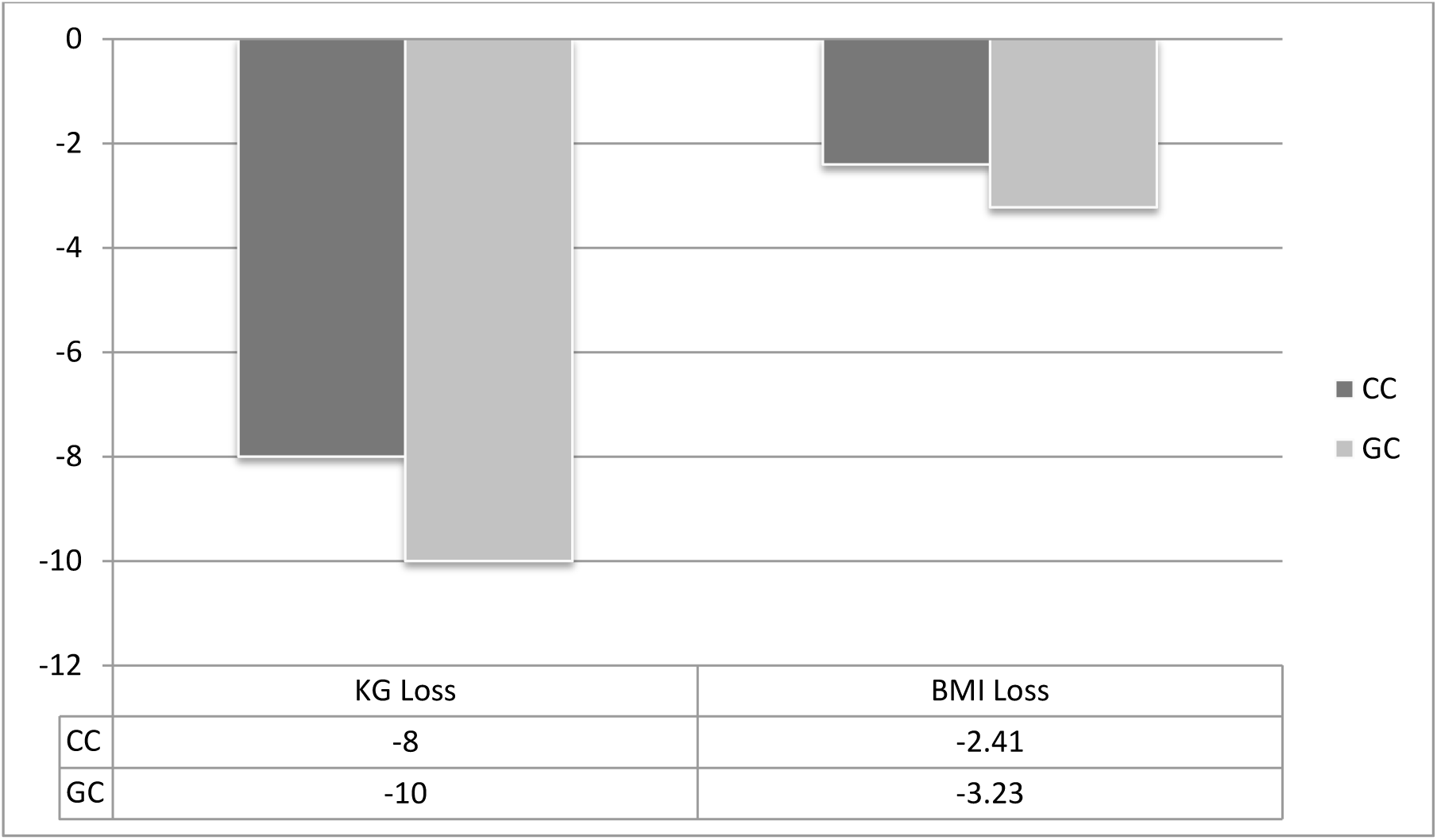
PPARG2 gene variants, weight and BMI change pre and post intervention.

### FTO

average weight loss for T (n=12) was −10kg with a BMI loss of −3.04, vales were significantly different pre and post intervention (t (22) = 4.61, P < 0.05). A allele (n=4) average weight loss was −9kg and BMI loss of −2.65 and values were significantly different (t (6) = 2.49, p = 0.02). TA individuals (n=22), BMI loss was −2.3 and weight loss was −7kg and significant differences (t (42) = 4.76, p < 0.05). Between groups, there was no significant different between T homozygous and A homozygous (t (14) = 7.79, P >0.05) or for T homozygous and TA heterozygous (t (32) = 3.59, P > 0.05). However a significant effect was found between A homozygous and TA heterozygous for BMI values (t (24) = 3.04, p = 0.002).

**Graph 4:**
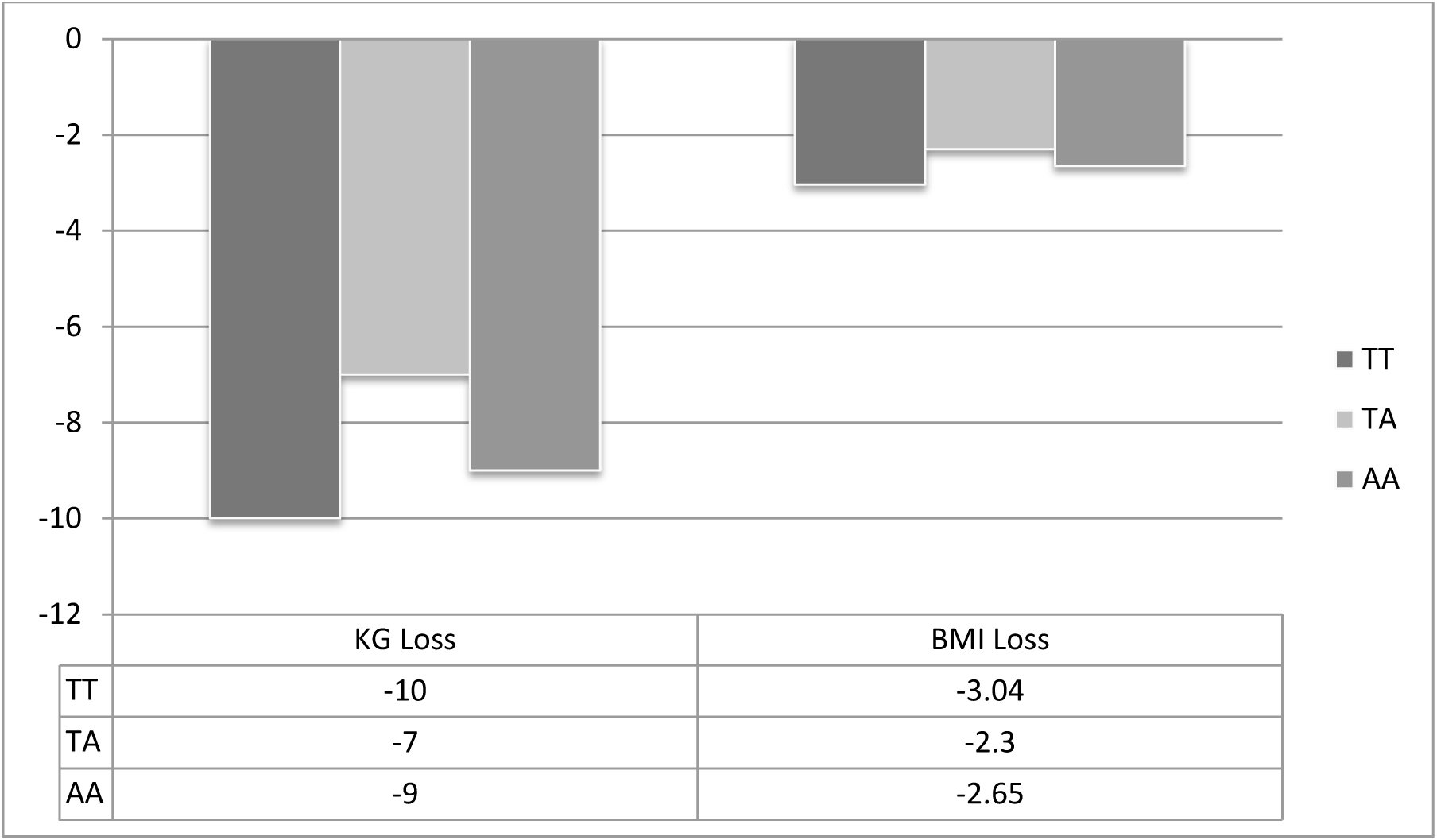
FTO gene variants, weight and BMI change pre and post intervention.

## Discussion

The field of obesity research is expanding to find a more complex aetiology surrounding the disease. New research suggest that adipose tissues now serves as an endocrine system, and secretes adipocytokines including adioponectin, IL-6, TNF-a and leptin. These cytokines are involved in systemic insulin responses and play a role in increasing symptoms of obesity and potentially leading to type-2 diabetes ^(4)^.

### TFAP2B – rs987237

The activating enhancer binding protein (AP)-2b transcription factor gene (**TFAP2B**) has been identified as a susceptible gene to increasing BMI and weight gain through its regulation of adipokine gene expression.

A previous study outlined the effected of the TFAP2B polymorphisms and found that the specific polymorphisms rs987237 had an overexpression of TNF-a in adipose cells ^(5)^. TNF-a produced in adipose cells has been implicated in systemic insulin resistance which in turn has been one of the major contributors to the pathogenesis of obesity ^(6)^. The expression of the risk allele “A” has been associated with decreased adiponectin levels in subcutaneous tissue.

Individuals with one or two of the G alleles have been shown to have significantly less leptin ^(8)^. Leptin is a hormone predominantly made by adipose cells that regulates energy balance by inhibiting hunger levels in the body. A lower level of circuiting leptin doesn’t inhibit hunger effectively; the individual doesn’t feel satiated from eating. This could potentially explain why carriers of the risk allele are known to have increased weight compared to their counterparts. Decreasing leptin concentrations are responsible for the starvation-induced suppression of the hypothalamic-pituitary-gonadal axis and can lead to insulin resistance, altering the metabolism of carbohydrate intake ^(9)^.

### FTO – rs993609

The rs993609 variant of the FTO gene is one of the most common genotypes that has known influence to the response of weight reduction therapy. The A allele of this gene has been shown to be the risk allele that is associated with lower insulin sensitivity and increased BMI levels ^(10)^. A previous study identified the FTO rs9939609 can interact with the macronutrient composition in individuals. The risk allele carriers (A allele) responded better to a high fat and low carbohydrate diet, but had higher drop-out rates from the study. High drop-out rates could be explained by a study in 3000+ individuals showing that the AA genotypes had a greater appetite compared to other genotypes, which suggests that this risk allele is likely to influence appetite and reduce satiation ^(11)^. The Homozygote T allele individuals had smaller decreases in resting energy expenditure (REE) and insulin release on the low fat higher carbohydrate diet ^(12)^.

In the present study, A allele individuals (both homozygous and heterozygous) started with the highest weight of all the participants, which could be explained by increased appetite levels and less satiation from their food intake. These individuals should preferentially modify caloric intake through more fat in place of carbohydrates. However this equates to a lower amount of total food consumed which could be difficult to adhere to long term due to the small satiating effects of the rick allele. A recent study also outlined potential differences in response to protein intake with the FTO *rs993609* and although protein intake was not changed during this study, normal day to day differences in intake could have affected an individual’s response ^(13)^.

### PPARG2 rs1801282

The peroxisome proliferator-activated receptor gamma 2 (PPARG2) gene is an extensively studied gene in relation to obesity. PPARG has been shown to play a crucial role in adipose tissue formation as well as controlling insulin. It is the insulin response by transcriptional stimulation of adipocyte-
genes which affects lipid storage, glucose and fatty acid uptake that is associated with metabolic disorders such as obesity ^(14)^.

Most studies outline the association between the polymorphism rs1801282 (Pro12Ala) and BMI. Carriers of the C allele have been shown to have naturally higher BMI when compared to non-carrier G or heterozygote G/C counterparts. A previous study assessed the interaction of macronutrient composition and BMI risk. They found a greater obesity risk in subjects who had a high carbohydrate intake with the C (Ala12) allele when compared to the non-risk G (Pro12Pro) allele. Further to this, individuals who had a high intake, combined with one just one C allele and one A allele of the FTO (rs9939609) gene had a further increased risk of obesity and higher BMI ^(15)^. The present study did not replicate significant results associating the PPARG Ala and FTO A allele for weight pre intervention (where individuals were consuming higher levels of CHO, (T (31) = 0.54, p = 0.29). However individual data showed these risk allele individuals did have higher weight and BMI pre intervention; average weight 90.4kg and BMI 28.1 kg/m^2^, compared to non-risk; average weight 89.4kg and BMI 27.6kg/m^2^. Overall values in this study found that homozygote G variant of the rs1801282 lost the least amount of weight compared to their counterparts. The reasoning behind this may be due to macronutrient split. C allele individuals have higher weight on a high CHO diet, suggesting their split should favour lower CHO and higher fat. This enhanced obesity risk on a high CHO diet can be explained that in the C allele individuals, CHO cannot activate the PPARG protein, which worsens receptor activity and decreases metabolic health. In G allele individuals, their response to CHO is better for weight loss, however, this variant also is responsible for the increased storage of fatty acids in adipose tissue on a high unsaturated fat diet ^(16)^. This explains why in this study, although G allele carriers still lost weight and BMI, it was one of the lowest values of all. These individuals should be cautious of over consuming on unsaturated fats and ensure a balanced diet whilst in a calorie deficit.

## TCF7l2 rs7903146

### The transcription factor 7-like 2 (TCF7L2)

Transcription factor 7-like 2 (TCF7L2) is involved in several functions within the cell. Its main involvement is in Wnt signalling pathways in various tissues within the body including fat, liver, brain and muscle cells. The Wnt signalling pathway is a strong activator of mitochondrial biogenesis which in turn leads to increased production of reactive oxygen species; inflammatory molecules which can damage cells and tissues. TCF7L2 regulates glucose metabolism in liver and pancreatic tissues and polymorphisms in this gene have clinical significance in type 2 diabetes and gestational diabetes through decreased insulin production, B-cell dysfunction and altered hepatic gluconeogenesis ^(17)^.

The current study shows similar findings to previous data ^(18)^. Results showed that the C variant of the genotype had greatest weight loss compared to the other variants. Individuals in this group responded better to the higher fat content of the diet after swapping out carbohydrates. This is not the case for the homozygous T or the heterozygous C/T. The C variant provided a protective mechanism by which the individuals still lost weight, despite being on a higher fat diet. However individually the T carriers did not lose a significant amount of weight pre and post intervention.

These results suggest that the TCF7L2 rs7903146 may operate via impaired glucagon-like peptide 1 secretion, which is stimulated by fats rather than carbohydrates ^(19)^. Carriers of the risk allele (T) in this gene need a moderate carbohydrate intake in order to lose any fat mass as lipid levels are regulated in an insulin dependent manner.

## Conclusion

Gene-diet interactions have paved a way for new scientific discoveries surrounding chronic disease related conditions, but also for a more personalised view of weight loss. To support a dietary intervention for weight loss, considering a person’s genetic predisposition, genetic variants determining dietary intake and metabolism and their genetic variants to exercise are all valuable tools to ensure advice is individually tailored.

The outcome of this study shows a need to control sugar intake in the general athletic population. Although everyone was in a calorie deficit for the study, the specific macronutrient swaps that were made impacted on the ability to lose body weight over the intervention period. Data from this study has outlined specific macronutrient interactions with single nucleotide polymorphisms in multiple genes and ultimately suggests a need for more personalised nutrition.

Personalised diet preparation that includes DNA analysis may prove to be more beneficial to those wishing to lose weight and fight obesity than preparation without. Weight loss that is efficient, effective and most importantly sustainable is key, with genetic testing used in conjunction with standard analysis helping enable this.

## Appendix

### Overall Averages from all data – minus outliers

|  | WeightPre | WeightPost | Height | kCAL | BMI Pre | BMI Post | Weight Loss over 24 WK | BMI Change over 24 WK |
| --- | --- | --- | --- | --- | --- | --- | --- | --- |
| Averages | 91 | 81 | 179 | 2548 | 28.1 | 25.2 | -9 | -2.9 |

t-Test: Paired Two Sample for Means between weight pre and postintervention

|  | Variable 1 | Variable 2 |
| --- | --- | --- |
| Mean | 90.55263158 | 81.21052632 |
| Variance | 22.41607397 | 30.49502134 |
| Observations | 38 | 38 |
| Pearson Correlation | 0.673551912 |  |
| Hypothesized Mean Difference | 0 |  |
| df | 37 |  |
| t Stat | 13.69 |  |
| P(T<=t) one-tail | 0.00 |  |
| t Critical one-tail | 1.69 |  |
| P(T<=t) two-tail | 0.00 |  |
| t Critical two-tail | 2.03 |  |

t-Test: Paired Two Sample for Means for BMI pre and post intervention

t-Test: Paired Two Sample for Means for BMI pre and post intervention
|  | *Variable 1* | *Variable 2* |
| --- | --- | --- |
| Mean | 28.14807544 | 25.24325077 |
| Variance | 2.956082802 | 3.544872318 |
| Observations | 38 | 38 |
| Pearson Correlation | 0.732555139 |  |
| Hypothesized Mean Difference | 0 |  |
| df | 37 |  |
| t Stat | 13.50 |  |
| P(T<=t) one-tail | 0.00 |  |
| t Critical one-tail | 1.69 |  |
| P(T<=t) two-tail | 0.00 |  |
| t Critical two-tail | 2.03 |  |

Data from 4 genes which found associations for

Data from 4 genes which found associations for
|  | TCF7L2 |  |  | TFAP2B |  |  | PPARG2 |  |  | FTO |  |  |
| --- | --- | --- | --- | --- | --- | --- | --- | --- | --- | --- | --- | --- |
|  | rs7903146 |  |  | rs987237 genotype |  |  | rs1801282 genotype |  |  | rs9939609 genotype |  |  |
|  | C | T | CT | A | G | GA | C | G | GC | T | A | TA |
| **N** | 24 | 9 | 5 | 21 | 1 | 16 | 31 | 1 | 6 | 12 | 4 | 22 |
| **Weight Pre (kg)** | 90 | 86 | 89 | 90 | 89 | 91 | 90 | 90 | 90 | 89 | 91 | 90 |
| **Weight Post (kg)** | 81 | 83 | 83 | 82 | 78 | 81 | 82 | 84 | 80 | 79 | 83 | 83 |
| **Height (cm)** | 179 | 179 | 179 | 170 | 177 | 179 | 179 | 182 | 180 | 180 | 179 | 180 |
| **KCAL** | 2558 | 2459 | 2494 | 2566 | 2450 | 2532 | 2553 | 2450 | 2550 | 2500 | 2543 | 2573 |
| **BMI Pre** | 27.96 | 26.39 | 27.85 | 27.76 | 28.41 | 28.19 | 28.01 | 27.17 | 27.8 | 27.55 | 28.57 | 28 |
| **BMI post** | 25.31 | 25.64 | 25.88 | 25.49 | 24.9 | 25.32 | 25.6 | 25.36 | 24.57 | 24.5 | 25.91 | 25.71 |
| **Weight Loss (kg)** | -8 | -2 | -6 | -7 | -11 | -9 | -8 | -6 | -10 | -10 | -9 | -7 |
| **BMI loss** | -2.6 | -0.75 | -1.9 | -2.27 | -3.51 | -2.87 | -2.41 | -1.81 | -3.23 | -3.04 | -2.65 | -2.3 |

